# A comparison of commercially available *Saccharomyces* mead yeasts

**DOI:** 10.64898/2026.02.27.708468

**Authors:** Bálint Németh, Zoltán Kállai, Aizhan Toxeitova, Gergő Horváth, Zsuzsa Antunovics, Andrea Harmath, University of Debrecen Biotechnology BSc class of 2026, Matthias Sipiczki, István Pócsi, Walter P. Pfliegler

**Affiliations:** Department of Molecular Biotechnology and Microbiology, Faculty of Science and Technology, University of Debrecen, Egyetem tér 1., H4032 Debrecen, Hungary; Doctoral School of Nutrition and Food Sciences, University of Debrecen, Egyetem tér 1. / Böszörményi út 138., Debrecen, H4032, Hungary; Institute of Horticulture, University of Debrecen, Debrecen, Böszörményi út 138., Debrecen, H4032, Hungary; Department of Genetics and Applied Microbiology, University of Debrecen, Egyetem tér 1., Debrecen, H4032, Hungary; Department of Medical Microbiology, Faculty of Medicine, University of Debrecen, Egyetem tér 1., Debrecen, H4032, Hungary; Doctoral School of Pharmaceutical Sciences, University of Debrecen, Nagyerdei krt. 98., Debrecen, H-4032, Hungary; Faculty of Science and Technology, University of Debrecen, Egyetem tér 1., Debrecen, H4032, Hungary; HUN-REN-UD Fungal Stress Biology Research Group, HUN-REN Hungarian Research Network, Debrecen, Hungary

**Author notes:** these authors contributed equally (Bálint Németh, Zoltán Kállai). list of group authors: Nikoletta Benyovszky, Domonkos Boldogh, Bence Boros, Janka Boros, Krisztián Boros, Liliána Dancsi, Máté Fejes, Lili Nóra Herédi, Anna Ágnes Kertész, Zsanett Madarász, Zsanett Mária Nagy, Hanna Barbara Pércsi, Petra Sárkány.

**Keywords:** Honey wine, mead, yeast, volatile compounds, commercial strains

## Abstract

We present a comparative analysis of 13 yeasts available for mead (honey wine) fermentation, a source of *Saccharomyces cerevisiae* diversity *t*hat has not yet been analyzed in detail. Using genomic, phenotyping, and analytic methods, we show that currently available mead yeasts belong to various clades of the species, most commonly to the Commercial Wine clade (5 of 13 samples). Mead yeasts in this group displayed genome structure variations and occasional loss of killer activity, despite being closely related. Historic European and traditional African mead isolates with sequenced genomes were found not to be closely related to any contemporary mead yeast product. The 13 yeasts tested here displayed high variability in oenological characteristics and in aroma production. Maximum ethanol tolerance ranged from 15 to 22% v/v, however, the most tolerant strain produced lower ethanol levels and retained high fructose content in experimental meads. The most abundant aroma components produced in meads were ethyl acetate, ethyl caprylate, isoamyl alcohol, and ethyl caprate, with similar aroma profiles in members of the Commercial Wine clade, and pronounced differences among other yeasts. Our results contribute to the knowledge of *Saccharomyces* yeasts in various fermentation environments, adding mead to the list of alcoholic beverages with a known diversity of starter cultures. Our results may aid strain selection for honey wine fermentations and inspire strain improvement.

## Introduction

Mead, or honey wine, is an alcoholic beverage made from bee honey and water with or without additional fruits, spices, and other flavoring components (Kawa-Rygielska et al., 2019). Mead has a historical and cultural importance as a beverage produced since millennia (Moe and Oeggl, 2013) with evidence on partially honey-based fermentations from the 7^th^ millennium B.C.E. from China, and palynological evidence suggesting meadowsweet-flavored and other types of mead from the 3^rd^ millennium to ∼400 B.C.E. from Northern- and Central Europe and the Caucasus (Kvavadze et al., 2006; Moe and Oeggl, 2013). Written accounts of mead production from Roman authors exist dating back to 37 B.C. (by Varro) and 70 C.E. (by Columella) (Forster and Heffner, 2001; McGovern, 2010).

Mead has recently gained interest both in food microbiology (Gaglio et al., 2017; Souza et al., 2024) and chemistry (Kawa-Rygielska et al., 2019; Pereira et al., 2019; Švecová et al., 2015; Webster et al., 2025) and in the home-brewing communities (Schramm K, 2003). The traditional production process often includes the heating or boiling of honey must before fermentation for protein removal and sterility. Alternatively, honey must is fermented without heat treatment (Bednarek and Szwengiel, 2020; Kahoun et al., 2017). Relatively few scientific accounts exist on indigenous mead fermentations, and T’ej (in Amharic, or Mess in Tigrina language), a ceremonial drink made in Ethiopia and Eritrea is one of the most well-known examples. This mead is flavored with gesho/geshu leaves (*Rhamnus prinoides*) (Dhyani A. et al., 2019).

Recent publications helped to gain insight into the organoleptic variations of mead (Senn et al., 2021; Simão et al., 2023; Švecová et al., 2015), into novel technological possibilities (Iglesias et al., 2014; Pereira et al., 2013; Ramalhosa et al., 2011), and also described the characterization of novelty products based on mead (Araújo et al., 2022). It is generally concluded that the yeast strain selection has a significant effect on the aroma profile of meads (Bednarek et al., 2019; Senn et al., 2021). Recently, some studies characterized various *Saccharomyces* (Bednarek et al., 2019; Schwarz et al., 2020) and non-*Saccharomyces* yeasts including *Kluyveromyces lactis, Pichia jadinii*, and *Torulaspora delbrueckii* (Van Mullem et al., 2022) for their applicability in mead fermentations. Meads fermented with lactic acid producing yeast species (Peepall et al., 2019) or the probiotic yeast (*S. cerevisiae* var. ‘*boulardii*’) have been described in detail (de Souza et al., 2024). However, most meads with information on the applied strain and in fact, most available recipes (Ratliff R. D., 2017; Schramm K, 2003; Senn et al., 2021) resort to using well-known wine and Champagne yeasts that are widely available as active dry yeast starter products. Senn et al. (2021) listed the following commercially available yeast strains as ones used in traditional American meads in a comparative study: Lalvin 71B, K1-V1116, BM 4×4, EC 1118, D21, D47, ICV D254, RC212 W15; Fermentis BC S103; Fermichamp Montrachet; Lallemand Abbaye, CBC-1; Munton’s Ale; Renaissance Yeast VIC-23; Safale US-05; Uvaferm GHM; White Labs WLP007, WLP775. However, in this case, various samples were fermented from different honey types, preventing comparisons on the yeasts’ contribution to the organoleptic characters of the meads. A few studies enabled comparisons of the yeasts’ effect on the final product: Pereira et al. (2009) compared yeasts isolated from honey in Portugal with one unspecified commercial and a laboratory *Saccharomyces*, while Pereira et al. (2019) described meads fermented with Lalvin’s QA23 and ICV D47. Avîrvarei et al. (2023) compared Lalvin DV10 and EC 1118 (in addition to a *Torulaspora* strain). Of the aforementioned yeast strains, all are widely used wine or Champagne yeasts except for Lallemand Abbaye, Munton’s Ale, Safale US-05, WLP007 (ale strains), CBC-1 (strain for bottle refermentation), and the commercially not available Portuguese honey isolates of Pereira et al. (2009).

There is very little data on *Saccharomyces* yeast isolates from indigenous and historic meads, constrained to whole genome sequences, without specific tests for aroma profiles and other characteristics in honey must fermentations. The strain DBVPG1853 from Ethiopia was sequenced by three studies (Hose et al., 2015; MacLean et al., 2017; Strope et al., 2015). Its source is given as “white tecc” honey wine, its collection date is unknown, but it was deposited before 1959. Three Ethiopian isolates from “white and red tecc” were included in a global analysis of Peter et al. (2018), namely DBVPG1841, DBVPG1843, and DBVPG1848 (with identifiers BPG, BPH, and BPI). These were also deposited in collection before 1959. Han et al. (2021) sequenced nine newly collected T’ej isolates and one isolate from geshu leaves from Ethiopia. Furthermore, DBVPG1853 was also recently sequenced by the National Collection of Yeast Cultures (NCYC, UK) in a BioProject (PRJEB42916) under the name NCYC 3313. This BioProject also included the following honey wine isolates: NCYC 356 (a mead production isolate from 1953, isolated from a culture of NCYC 93 after being used for mead production for 2 years by C. H. Ridge; note that NCYC 93 is a yeast of unknown origin from 1925); NCYC 357 (an isolate collected in 1951 from an “Avize-Cramant mead”, thus, a mead from the Côte des Blancs subregion of Champagne in France), and NCYC 358 (from the lees of an unspecified plum mead). Thus, of the thousands of sequenced yeasts genomes (Loegler et al., 2024), merely a small minority comes from honey and honey wine fermentations, while beer and wine starter cultures, including globally available commercial strains, are much better known. Furthermore, the genomes of the NCYC mead strains have not yet been analyzed phylogenomically.

Several companies have recently begun to commercialize specific mead yeasts as active dry or liquid fermentation starters, without specifications on their origin. In this study, our aim was to test and compare such yeasts and to analyze their phylogenomic positions. We obtained mead yeasts from retailers in Europe and isolated single-cell derived lineages from these products, and conducted phenotyping, genetic fingerprinting, and experimental honey must fermentations with them. We also performed whole genome sequencing and a phylogenomic analysis on the commercial strains, and compared their relationships with the previously sequenced honey and mead isolates. Our results may help commercial and home-brewers to evaluate mead yeast for fermentations. Our study on these yeasts was integrated into the Biotechnology BSc program’s General and Applied Microbiology courses at the University of Debrecen, thereby providing an example of applied research on yeasts integrated into regular university practical classes. Students passing the course are listed in this work as group authors.

## Materials and Methods

### Strains

We obtained yeast strains marketed as mead or honey wine strains after internet searches in Google and on various retailers’ websites. Altogether, for the purposes of our study, we managed to obtain 13 mead yeast strains from various retailers, covering all available mead yeasts at the time of purchase except for one strain only marketed in Argentina (The Beer Company: Turbo Hidromiel Dulce, Google search, 31.01.2024). Yeast strains obtained for this study originated from Czechia, Poland, Sweden, UK, and USA (Table 1.). Single-colony isolates were obtained after purchase on YPD medium, and yeasts were given identifiers (ranging from UDeb-MY0001 to 0013). Most of the mead yeasts were marketed as *Saccharomyces cerevisiae*, while some as the taxonomically obsolete *S. bayanus*. Eight samples were active dry yeast products, five were liquid yeasts (Table 1.). Single-cell derived subclone cultures were deposited in the collection of the Department of Molecular Biotechnology and Microbiology (University of Debrecen) as frozen stocks in yeast extract peptone dextrose medium (YPD) (2% glucose, 2% peptone, 1% yeast extract) supplemented with 30% v/v glycerol.

**Table 1.** Strains collected for this study.

| Isolate name in collection | Formulation | Product name | Country of origin |
| --- | --- | --- | --- |
| UDeb-MY0001 | active dry | Mangrove Jack mead M05 | USA |
| UDeb-MY0002 | active dry | Gozdawa mead | Poland |
| UDeb-MY0003 | active dry | Bulldog mead | UK |
| UDeb-MY0004 | active dry | Coobra mjöd | Sweden |
| UDeb-MY0005 | active dry | Enovini mead | unspecified |
| UDeb-MY0006 | active dry | Alcomat mead | Poland |
| UDeb-MY0007 | liquid yeast | Wyeast dry mead 4632 | USA |
| UDeb-MY0008 | liquid yeast | Wyeast sweet mead 4184 | USA |
| UDeb-MY0009 | liquid yeast | White Labs WLP720 wweet mead/wine | USA |
| UDeb-MY0010 | active dry | Starowar mead *S. bayanus* | Poland |
| UDeb-MY0011 | active dry | Strongferm mead *S. bayanus* | Poland |
| UDeb-MY0012 | liquid yeast | Dolské kvasinky medovina strong | Czechia |
| UDeb-MY0013 | liquid yeast | Dolské kvasinky medovina | Czechia |

### PCR-fingerprinting

Genomic DNA extraction was performed according to the protocol described by Hanna and Xiao (2006), and DNA concentrations were standardized to 100 ng/μL for all isolates. Genomic DNA samples were stored in 1× TE (Tris-EDTA) buffer at −20⍰°C. Multiplex fingerprinting was based on the use of the primer sets designed to amplify interdelta elements, microsatellites of the genes YLR177w, YOR267c, and the ITS (internal transcribed spacer) region as described by Imre et al. (2019). Gel electrophoresis was performed using 2% agarose TBE gels, at 100 V, for 90 minutes, with 4 µl of PCR product. Band patterns were visually compared after photographing gels with UV-transillumination.

### Karyotyping

Karyotyping was performed using 1 % agarose gel (chromosomal grade, Bio-Rad, Hercules, CA, USA) by a counter-clamped homogenous electric field electrophoresis device (CHEF-Mapper; Bio-Rad). The following running parameters were used: run time 26 h, voltage 6 V/cm, angle 120°, temperature 14 °C, and pulse parameters 60 to 120 s. As a control, a haploid *S. cerevisiae* control was used. After electrophoresis gels were stained with ethidium bromide and washed in sterile water for 48 h before photographing using UV-transillumination, subsequently, band patterns were compared.

### Phylogenomics

A reference-based mapping and called SNP-based phylogenomic approach was used to cluster the mead isolates into the described clades of the species. To enable comparisons with previously published clades, mosaic groups, and even strains to an unprecedented level, we relied on the Compendium of *Saccharomyces* genomes (Németh et al., under publication). The isolates were included in a large-scale phylogeny of nearly 5000 *S. cerevisiae* genomes. Their cladal origins were determined based on clustering with known members of clades described by Loegler et al. (2024). As the phylogeny included all previously published genomes available for commercial strains, we were also able to determine whether a given mead isolate showed stain-level identity to any previously described yeast.

The DNA isolated as described above was subjected to short-read sequencing either by Novogene (Germany) as a paid service or by the Core Facility of the University of Debrecen. In the latter case, library preparation was performed using tagmentation with the Nextera DNA Flex Library Prep kit (Illumina, San Diego, CA, USA) according to the manufacturer’s protocol. Sequencing was performed using 150 bp paired-end reads on an Illumina NextSeq 500 system, with approximately 50–80× coverage of the nuclear genome. In the case of Novogene, sequencing was performed using paired-end 150 bp technology on an Illumina NovaSeq X Plus system. During library preparation, genomic DNA was fragmented to an average size of 350 bp, and size selection was carried out using sample purification beads. Raw reads of new samples were deposited to NCBI SRA under BioProject PRJNA1420404. The Illumina FASTQ sequencing files were trimmed and filtered using fastp for further analysis (Chen et al., 2018). We mapped Illumina reads to a concatenated reference panel of eight *Saccharomyces* species (*S. arboricola, S. cerevisiae, S. eubayanus, S. jurei, S. kudriavzevii, S. mikatae, S. paradoxus*, and *S. uvarum* downloaded from GenBank) to avoid introgressed regions in several clades and non-*cerevisiae* subgenomes in hybrids skewing phylogenomic positions. Mapping was performed using the mem option of BWA 0.7.17. (Li and Durbin, 2010). Sorted BAM files were obtained using Samtools 1.7. (Danecek et al., 2021; Li et al., 2009) and Picard-tools 2.23.8. was used to mark duplicated reads (Poplin et al., 2018; Van der Auwera et al., 2013). Using BAM files, local realignment around indels and joint variant calling and filtering for the strains and isolates were performed with GATK 4.1.9.0. (Poplin et al., 2018; Van der Auwera et al., 2013) with regions annotated in the S288c reference as centromeric regions, telomeric regions, or LTRs excluded. First, genomic VCF files were obtained with the Haplotype Caller for the *S. cerevisiae* chromosomes of the concatenated reference, and joint genotyping of the gVCF files was applied. Following the joint calling, only SNPs or only indels (insertions and deletions) were selected in the resulting VCF files. SNPs were filtered according to the parameters as follows: QD < 5.0; QUAL < 30.0; SOR > 3.0; FS > 60.0; MQ < 40.0; MQRankSum < -12.5; ReadPosRankSum < - 8.0. INDELS were filtered according to the parameters QD < 5.0; QUAL < 30.0; FS > 60.0; ReadPosRankSum < -20.0 (Fay et al., 2019). Indels were then left-aligned. For the final VCF files, indels and SNPs were merged, filtered and non-variant sites were removed. The multisample.vcf was converted to.gds file with the snpgdsVCF2GDS function of SNPrelate (Zheng et al., 2012). Then, a dissimilarity matrix was created using the snpgdsDiss function of SNPrelate. The matrix was imported into the ape 5.8-1 package (Paradis and Schliep, 2019), creating a neighbor-joining tree and iTOL was used to visualize the output (Letunic and Bork, 2019). Clades and clade colors were taken from Loegler et al. (2024).

### Comparative genomics

For allele frequency analysis, variants in the individual strains were selected from the above-mentioned 8-species mapping cohort.vcf file and exported to a.csv file using the query option of SAMtools/BCFtools 1.10.2. (Danecek et al., 2021; Li et al., 2009). Allele frequency plots were obtained from these using a custom pipeline in R (R Core Team, 2021) using a for loop to calculate allele depth ratios from the exported.csv (defined as the ratio of each allele’s sequencing depth to the sum of all alleles’ depth at a given locus) for every genome and the ggplot2 package (Wickham, 2016) for data visualization. Depth ratio values for all alleles were plotted for each variant locus along the chromosomes. Allele frequencies were used to estimate ploidy, with the assumption that disomic chromosomes have allele ratios of approx. 1:0 or 1:1, trisomic of 1:0, 1:2, and 2:1, tetrasomic of 1:0, 1:3, 1:1, or 3:1, etc. (Rácz et al., 2025; Zhu et al., 2016). The results obtained from allele ratios were compared to coverage plots to determine whether any of the isolates are non-diploids. For coverage plots, reads were first mapped to the 8-species reference as described above and then, to the *S. cerevisiae* genome only after the non-hybrid nature of the given samples was determined. We used BEDTools 2.30.0 (Quinlan and Hall, 2010) to create per-base BedGraph files and then to calculate the median coverage of chromosomes in 10000 base windows sliding every 5000 bases. Plots generated from this data were corrected for ploidy and were used to identify potential aneuploidies. Rows of homozygosity (ROH) were identified using BCFTools 1.10.2. and visualized together with allele plots.

### Colony morphology on agar medium, invasivity, and flocculation in liquid medium

Strains were plated on YPD agar medium from OD_600_=1 suspensions in distilled water and incubated at 22°C for 10 days. Then, colonies were observed visually and photographed for their shapes, sizes, and colors. Colonies were washed off with tap water from the plates and any invasive growth into the agar was photographed. Strains were inoculated in 5 mL YPD liquid medium and incubated without shaking at 22°C for 10 days in test tubes to observe whether flocculation occurs.

### Sporulation test

In order to sporulate mead yeasts, potassium acetate agar was used (1% potassium acetate, 0.1% yeast extract, 0.05% glucose, 2% agar). Cultures were incubated on these plates for 10 days at 22°C, and sporulation was checked under 400× magnification using a light microscope.

### Extracellular protease and phospholipase activity

Strains pre-cultured on YPD agar were inoculated in 5 μL spots (5×10^4^ cells) onto protease and lipase test media and incubated for 7 days at 22°C. Protease secretion was followed on milk agar plates (0.3% w/v bouillon, 1% w/v peptone, 5% w/v milk powder, 2% w/v agar). Lipase secretion was tested on Tween agar plates (1% w/v peptone, 0.1% w/v CaCl_2_, 0.5% w/v NaCl, 1% v/v Tween, 2% w/v agar). After incubation, zones indicating enzyme production were observed.

### Spot plate assay stress and starvation phenotyping

All tests were conducted using a standard synthetic dextrose (SD) medium (0.17% w/v Yeast Nitrogen Base without amino acids, 0.5% w/v ammonium sulphate, and 2% w/v agar). Yeast suspensions of 0.5 MacFarland units were prepared using a densitometer, and a 10× dilution series was prepared in distilled water. From the series, 5 µl drops were plated onto the agar media, containing ∼10,000, ∼1,000, ∼100, ∼10, and ∼1 cells. Plates were incubated for 3 d and 10 d before visual evaluation. Growth was scored according to the spots showing yeasts growth. For high temperature tolerance, plates were incubated at 22°C, 37°C, and 39°C for 3 d. For starvation tests, synthetic low glucose (SLG), synthetic low ammonium (SLAD) with various ammonium sulfate concentrations (0.0066 g/l amounting to the standard SLAD recipe, 0.25 g/l, and 0.5 g/l amounting to the 5% and 10% value of the ammonium sulfate content of standard SD, respectively) were used. SD plates containing all vitamins and nucleotides in the standard composition except biotin and thiamin were also prepared, to these, small amounts of these vitamins were added, amounting to the 5% of the amount found in standard SD medium. A growth score was determined based on the number of spots showing clear growth.

### Stress phenotyping in liquid medium

Yeasts (OD_600_=1 suspension, 1 ml) were first incubated at 22°C in 50 ml of high-sugar SD liquid medium (350 g/l glucose) without shaking, with 1.2 g/l commercial yeast nutrient (Brewline, Canéjan, France) addition divided into 4 rations, after 1 d, 2 d, 3 d, and 7 d following inoculation (Ratliff R. D., 2017). For high osmotic stress tests, 100 µl of the pre-culture was inoculated into 5 ml high-sugar liquid SD media with 30%, 40%, 50%, and 60% glucose (w/v) concentrations. They were monitored for two weeks for growth and fermentation at 22°C. Results were scored after visual evaluation of yeast growth and fermentation. For alcohol tolerance tests, 100 µl of the pre-cultures was inoculated into liquid SD medium with various concentrations of ethanol, ranging from 14 to 22% v/v. After incubation at 22°C for 7 d, the media were briefly vortexed and samples were taken. A 10× dilution series was prepared from the samples and the above-described spot plate method was applied. Yeast growth was scored after 5 days of incubation.

### Killer activity and sensitivity tests

Production of the killer toxin and sensitivity of mead yeasts were evaluated using two types of killer yeasts: killer yeast 1 (K1, NCYC 232) and killer yeast 2 (K2, NCYC 738). To test the killer activity, suspensions of killer controls (20 μL, OD_600_=0.1) and mead yeasts were inoculated onto the YPD agar plates (OD_600_=1 suspensions, 10 µl), containing 0.003 g/L methylene blue, in which citrate-phosphate buffer was used to buffer the media to pH 4.5. These agar plates were kept at 22°C for a 3-day incubation period. After that, suspensions of pre-cultured sensitive yeast strains (NCYC 1006 1.0 McFarland, 100 μL, ∼200,000 CFU) were sprayed on the plates and incubated again for 4 days at 22°C. Sensitivity to killer activity of mead yeasts by the control sensitive yeast was observed by the appearance a halo that shows a decline of growth with a dark blue zone of dead cells around the edge. Sensitivity test: the two killer control yeasts were tested against all of the 13 mead yeasts. K1 and K2 killer yeasts were inoculated onto plates and mead yeasts were sprayed 3 days later around these, using the same setup used to test killer activity of the mead yeasts and incubated again for 4 days at 22°C. Halo and colony diameters for all killer tests were measured and their ratio was calculated in triplicates.

### Experimental fermentations

The mead yeasts were tested in small-scale experimental fermentations in triplicates, modeling homebrewing technics. Floral honey from the Hortobágy region, Hungary was obtained from a small-scale local producer, produced in the late summer of 2023. A single batch of honey was used and all 39 fermentations with the 13 mead yeasts were conducted at the same time. 300 g of honey was dissolved without heating in commercially obtained mineral water (HCO_3_^−^ 385 mg/l, Ca^2+^ 59 mg/l, Na^+^ 30 mg/l, Mg^2+^ 28.7 g/l). The pH of the honey must was 6.20, its original specific gravity was 1.095. The honey must was photographed before a white background in a clear glass with a diameter of 5 cm. Mead yeasts precultured on YPD plates were inoculated into the honey musts in 1 l glass flasks (3 ml, OD_600_=1). Fermentation was carried out at 18°C with vigorous shaking of the flasks twice every day for one week, and without shaking for 3 more weeks. The Tailored Organic Staggered Nutrient Additions (TOSNA 3.0) (Ratliff R. D., 2017; Schramm K, 2003) method was applied that is recommended for homebrewers: 1.2 g/l commercial yeast nutrient (Brewline) was added to fermentations, divided into 4 equal rations, 1 d, 2 d, 3 d, and 7 d following inoculation. After 4 weeks, meads were racked, and 1 dl samples without yeasts lees were used for analytics. Meads were also photographed before white background, and were visually checked for haziness.

### Oenological analytics

Fourteen oenological parameters of the triplicate mead samples were determined according to the official standards of the International Organisation of Vine and Wine (OIV, 2010) using a Lyza 5000 Wine FTIR analyzer (Anton Paar, Graz, Austria). These were ethyl-alcohol concentration, titratable acidity to pH 7.0 and to 8.2, pH; glucose, fructose, total volatile acid, malic acid, tartaric acid, glycerol, gluconic acid, total polyphenol, and extract contents; and density (Rakonczás et al., 2023).

### Semiquantitative analysis of volatile compounds

The volatiles’ profiling of the samples was carried out with a Bruker Scion 436 Gas chromatograph coupled with Bruker SQ mass spectrometer. The system was fitted with a DB-5MS capillary column (25 m 0.25 mm i. d. 1.0 µm film thickness). The carrier gas was helium 5.0, flow rate was 0,9 mL/min in constant flow mode. Samples were incubated in headspace vials at 40°C for 20 minutes in thermostat with no agitation. 1000 µL headspace sample was injected into the column. The transfer line was maintained at 230°C, the injector temperature was 250°C (20:1 split ratio). The oven was held at an initial temperature of 40°C for 2 minutes, then increased to 280°C at 10°C/min, and held at this temperature for 3 minutes. The mass spectrometer was operated in electron impact ionization mode (70eV) source temperature: 180°C; scanning rate: 1 scan s^−1^ and the mass spectra were recorded in full scan mode. Identification of the volatile compounds was based on mass spectrometric data obtained from the National Institute of Standards and Technology (NIST) (Version 2005) mass spectral library. As standards of the various volatiles were not measured, we expressed areas of the measured components as fractions of the highest area value, thereby, these semiquantitative measurements reflect the relative aroma concentrations in all of the experimental meads.

### Sensory evaluation

After the end of the lab course, a tasting event was organized where students could taste the experimental meads. The triplicate samples of each mead were first mixed, and a scoring sheet was prepared (FigShare https://doi.org/10.6084/m9.figshare.31375720.v1). Since the evaluation was not performed by trained mead (or wine) judges, the results are just briefly mentioned here and scoring results are uploaded to FigShare, noting that a trained panel may come to different results during a sensory evaluation.

### Statistical analysis

Statistical analysis was conducted using Statistics Kingdom (https://www.statskingdom.com/index.html). To compare more than two datasets at once, ANOVA (normal distribution), followed by Tukey HSD test or Kruskal-Wallis (non-normal distribution), followed by Dunn post-test (corrected for Benjamini-Hochberg FDR) was used to determine which datasets differed significantly. Principal components analysis (PCA) was applied to measured metabolites (for the means of all detected components’ values of the triplicate mead samples).

## Results

### Commercial mead yeasts and organizing the BSc students’ microbiology laboratory practice

We established a strain collection representing 13 strains of mead yeasts from 11 companies (Table 1.). Among these, 12 were explicitly marketed as yeasts for mead-making, while UDeb-MY0009 was a “sweet mead/wine” yeast. Five were sold as liquid starter cultures, eight as active dry yeasts. None of the yeasts had information on the origin of the strain (e.g. whether they were collected from traditional mead fermentations or not). UDeb-MY0010 and UDeb-MY0011 were marketed under the taxonomically obsolete name *S. bayanus*. In the results below, the yeast samples’ names are abbreviated by omitting the first part of the name referring to the University of Debrecen.

The yeasts were isolated with students during the General and Applied Microbiology lab course at our University in the spring semester of 2024. Students performed most of the phenotyping experiments (spot plates, killer tests, sporulation, colony morphology and invasivity, flocculation) and presented lab reports individually, with data visualization and statistics using online sources. The course ended in a written test focusing on general aspects of the microbiological work. Students passing the course successfully were included in the manuscript as group authors, and the manuscript, the submission to BioRxiv and to a journal were discussed with them during a special seminar. Written consent in order to be included in the manuscript was obtained.

### PCR fingerprinting and whole-genome phylogeny

The mead yeasts were subjected to DNA-isolation and genetic fingerprinting using a multiplex fingerprinting method. Electrophoretic band patterns were evaluated by the students after they performed the PCR, and several identical band patterns were observed (Figure 1.), suggesting possible strain-level identity of some samples. MY0003, MY0005, MY0007, MY0010, and MY0011 were identical with this analysis, despite originating from different companies in various countries (Table 1.).

**Figure 1.**
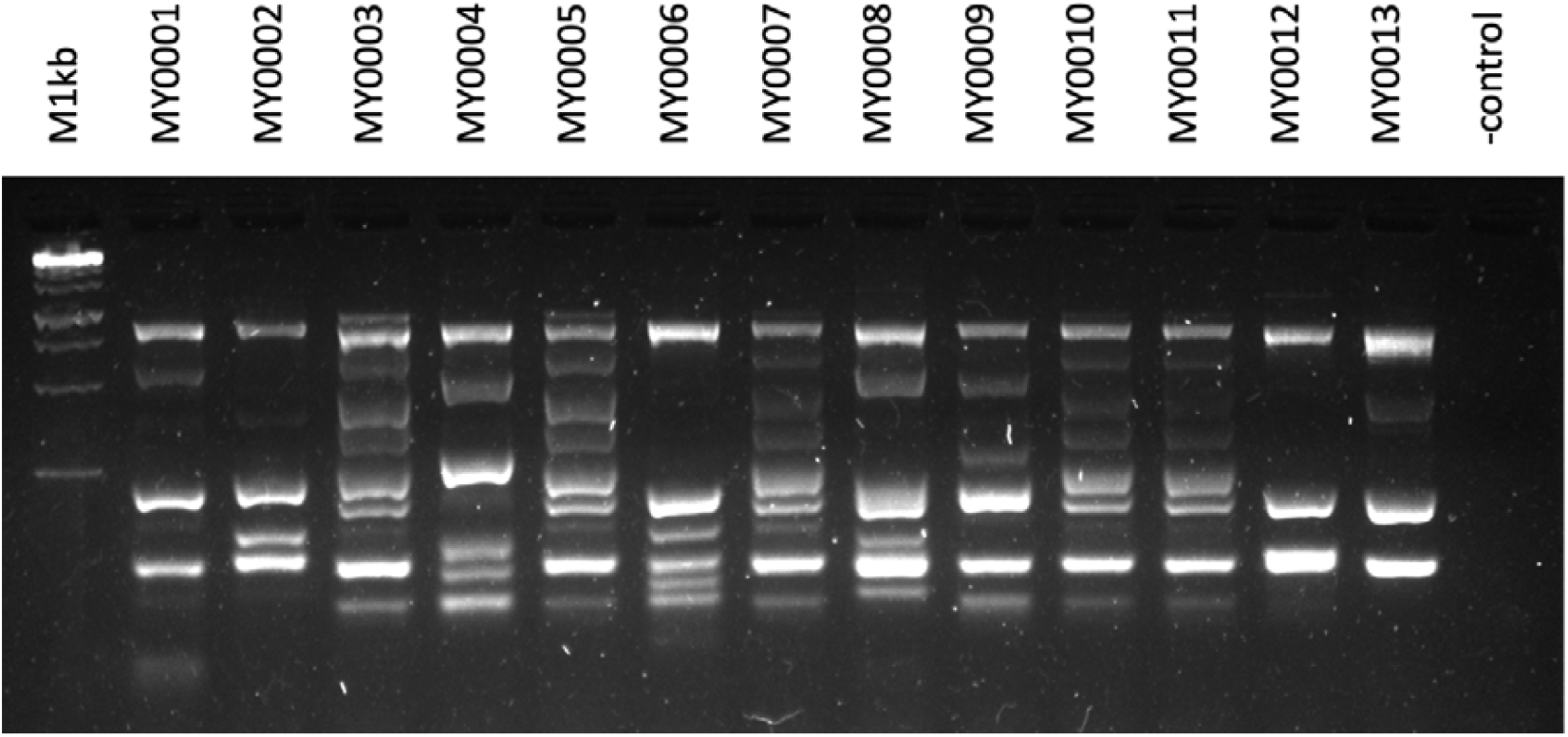
Multiplex PCR fingerprinting results. M1kb: 1 kilobase size marker. Mead yeast band patterns are followed by a negative control lane.

The identity of several samples with PCR-fingerprinting prompted us to perform a more detailed analysis involving whole-genome shotgun sequencing and phylogenomics to evaluate differences among samples on a finer scale, and to determine which clades they belong to. This bioinformatic analysis was done outside of the curriculum of the laboratory class. We compared the mead yeasts to ∼5000 *S. cerevisiae* genomes and identified which clades they belong to, and determined that none of them were hybrids. Thus, the *S. bayanus* samples proved to be simple wine yeasts of the species *S. cerevisiae*. One sample, MY0004 clustered with the Huangjiu or MAN2 clade of Loegler et al. (2024), with close relatedness to the commercial strain Ethanol Red. Two samples (MY0006, MY0009) belonged to the Mixed Origins 1 clade (MIX1), being related to a wide variety of baking strains and various other samples. MY0009 clustered closely to the WhiskyPure strain. MY0008 clustered in the UK Beer clade (UKB), with very close relatedness to the various samples of Safale S05 sequenced by previous studies. MY0001 belonged to the Australia Wine 3 clade (AUW3), being related to the commercial strain WLP099. MY0012 belonged to the Western Europe Wine clade (WEW). The Australia Wine 1 clade (AUW1) included MY0002 and MY0013. The aforementioned group of five identical strains in the fingerprinting analysis all clustered into the Commercial Wine (COW) clade, with very close relatedness to the commercial strain EC 1118, CBC-1, Fermicru 4F9, WY4766, and to each other (Table 2., Figure 2.). Regarding previously sequenced mead yeasts, the strain DBVPG1853 from Ethiopia and its four samples sequenced by four studies, along with ETPF1, ETPF2, ETPF3, ETPF4, ETPF5, ETPF6, ETPN8, and ETPF9 (Han et al., 2021) and BPI of Peter et al. (2018) clustered into the African Beer clade (AFB), as shown previously (Loegler et al., 2024). However, two samples from Ethiopia belonged to the Australia Wine 1 clade, namely BPG and BPH. The Mixed Origins 1 clade also included two Ethiopian samples, ETPF4 and ETPF5, with close relatedness to commonly available active dry baker’s yeasts. The historical mead yeast samples NCYC 357 and NCYC 358 clustered into the Georgia Wine clade (GEW), while NCYC 356 belonged to the Mixed Origins 1 clade.

**Table 2.** Details on the genomes of the 13 mead yeasts used in this study.

| Name | Clade | Ploidy | Hetero-zygosity | Aneuploidies | Plasmid | mtDNA | Coverage |
| --- | --- | --- | --- | --- | --- | --- | --- |
| MY0004 | Huangjiu | 2 | 0.0572% | no | no | yes | 65x |
| MY0006 | Mixed Origins 1 | 2 | 0.2923% | no | yes | yes | 52x |
| MY0009 | Mixed Origins 1 | 2 | 0.3539% | no | yes | yes | 81x |
| MY0008 | UK Beer | 4 | 0.4395% | +1*IV;<br>-1*VIII | yes | yes | 89x |
| MY0001 | Australia Wine 3 | 2 | 0.0621% | no | yes | yes | 77x |
| MY0012 | Western Europe Wine | 2 | 0.0118% | +1*IX | yes | yes | 157x |
| MY0002 | Australia Wine 1 | 2 | 0.0479% | no | yes | yes | 183x |
| MY0013 | Australia Wine 1 | 2 | 0.0121% | +1*I | yes | yes | 114x |
| MY0003 | Commercial Wine | 2 | 0.1348% | +1*XII | yes | yes | 49x |
| MY0005 | Commercial Wine | 2 | 0.1285% | no | yes | yes | 55x |
| MY0007 | Commercial Wine | 2 | 0.1590% | no | yes | yes | 132x |
| MY0010 | Commercial Wine | 2 | 0.1689% | no | yes | yes | 90x |
| MY0011 | Commercial Wine | 4 | 0.1676% | no | yes | yes | 136x |

**Figure 2.**
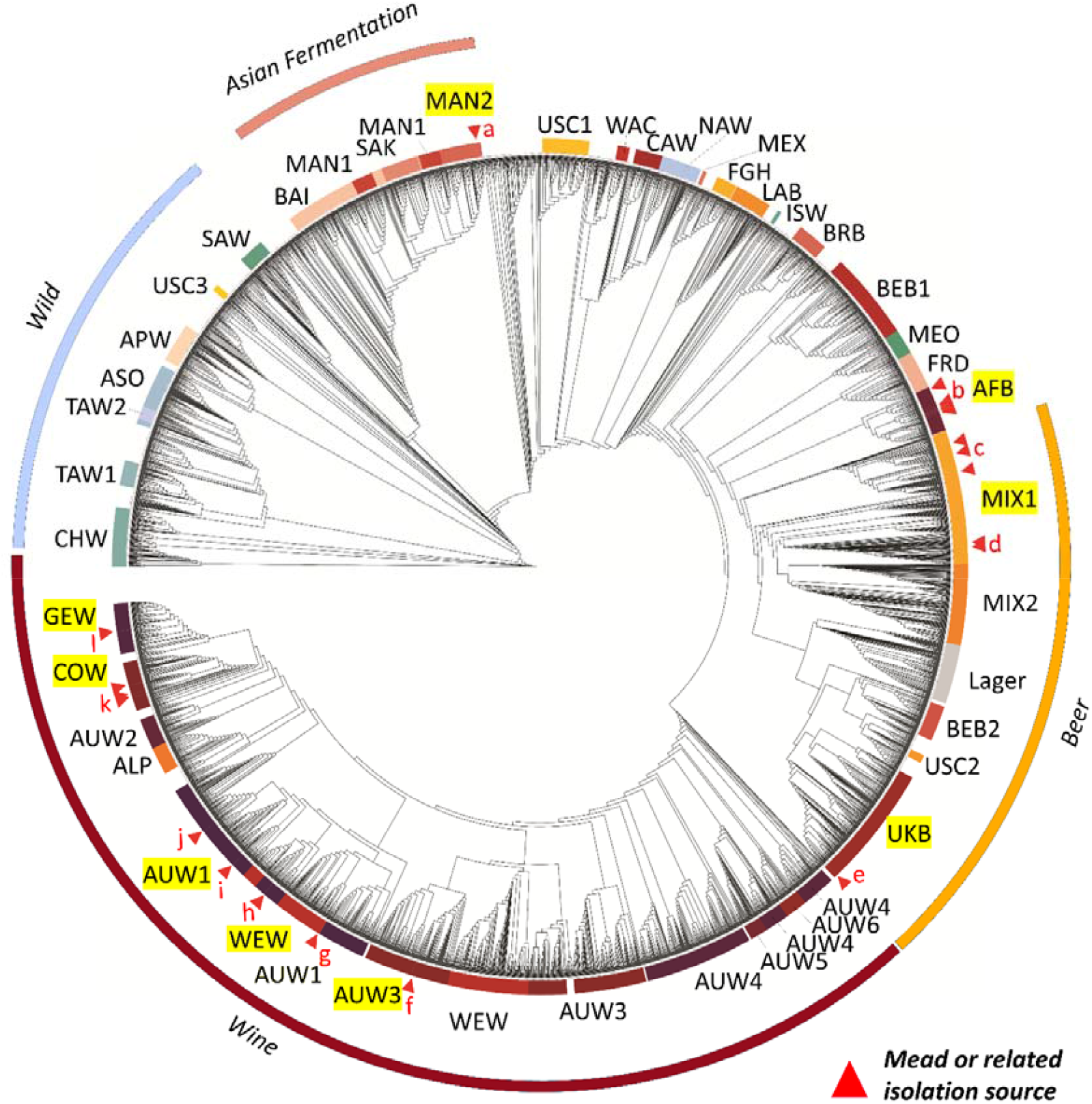
Phylogenomic dendrogram highlighting the placement of mead yeasts. Superclades as defined by Loegler et al. (2024) are marked as outer circles, clades are shown with color codes as inner circles along the dendrogram. Clades with mead yeasts are highlighted with yellow background. Positions of mead yeasts are shown with red triangles, marked with lower-case letters from a to l. Clade abbreviations: MAN2: Huangjiu. AFB: African Beer. MIX1: Mixed Origins 1. UKB: United Kingdom Beer. AUW3: Australia Wine 3. AUW1: Australia Wine 1. COW: Commercial Wine. GEW: Georgia Wine. Mead yeasts are marked with small-case letter as follows, in the order they appear on the phylogenomic tree: a (MY0004); b (ETPF2, ETPF6, ETPN8, ETPF1, ETPF11, ETPF13, ETPF3, ETPF9, ETPF, BPI, DBVPG1853, DBVPG1853Maclean, NCYC 3313, SACE_YDB); c (MY0006, NCYC 356, MY0009); d (ETPF4, ETPF5); e (MY0008); f (MY0001); g (MY0012); h (BPG, BPH); i (MY0013); j (MY0002); k (MY0003, MY0005, MY0007, MY0010, MY0011); l (NCYC 357, NCYC 358).

After establishing the phylogenomic placement of the mead yeast samples, we performed comparative genomic analyses and determined their ploidy, overall heterozygosity, and identified aneuploidies and runs of homozygosity, along with recording whether 2µ plasmid and mitochondrial DNA presence was inferred from the sequence coverage data (Table 2.). Data throughout the Results section is presented with the mead yeasts ordered according to clades below.

All mead yeast samples showed the presence of mitochondrial DNA, and all expect the Huangjiu clade MY0004 possessed a 2µ plasmid. The samples were mostly diploid euploids. MY0009 of the UK Beer clade was a tetraploid aneuploid yeast, while the allele ratio data of MY0011 of the Commercial Wine clade was consistent with a tetraploid euploid genome (Supplementary File S1.). In this clade, MY0003 also showed trisomy of chr. XII. MY0012 of the Western Europe Wine clade and MY0013 of the Australia Wine 1 clade showed trisomy of chr. IX and I, respectively (Table 2., Supplementary File S1.). Heterozygosity was the lowest, ∼0.012% in MY0012 and MY0013, which samples also showed characteristically simple band patterns during PCR-fingerprinting (Figure 1.). Other yeasts of the various wine clades showed heterozygosities ranging from 0.048% to 0.169%, the latter observed in the case of MY0010. Mixed Origins 1 mead yeasts were more heterozygous (0.292 to 0.354%), while the UK Beer MY0008 showed the highest value, 0.44%. The Huangjiu clade MY0004 showed low level of heterozygosity (0.057%). Runs of homozygosity (ROH) regions were also compared (Supplementary File S1.) and showed various patterns among the mead yeasts. Based on overall heterozygosity and the ROH calling, MY0012 and MY0013 can be considered homozygous genomes, while in other cases, the genomes clearly display heterozygous regions, even in the low-heterozygosity MY0002 genome (Supplementary File S1.). Among the five closely related mead yeasts of the Commercial Wine clade, ROH patterns were nearly, but not completely identical between MY0003 and MY0005, differing in the presence of a terminal ROH on chr. VII and in the presence of a short terminal ROH on chr. XIV in MY0005 that were not found in MY0003. MY0007 had similar patterns of ROHs with the same start and end points as MY0003, but lacked a terminal ROH on chr. XIII and one on chr. XV. MY0010 an MY0011 were almost identical in ROH patterns except for a terminal ROH on chr. XVI, and characteristically, they had a shared terminal ROH on chr. IV which was shorter than the terminal ROH of the other samples covering more than half of the chromosome. Thus, ROHs and aneuploidies were unique in all five samples of the Commercial Wine clade (Supplementary File S1.). We further characterized the genomes using electrokaryotyping. Occasionally, extra bands were observed, as in the two samples of the Mixed Origins 1 clade (MY0006, MY0009), in the Australia Wine 1 clade (MY0002, MY0013), and in the Commercial Wine clade (Figure 3.). In the latter, all samples showed an extra band in the region of the smallest chromosomes, and all expect MY0003 showed an extra band in the region between chr. XV – VII and IV. Among the five samples, MY0007, MY0010, and MY0011 showed identical karyotypes (Figure 3.).

**Figure 3.**
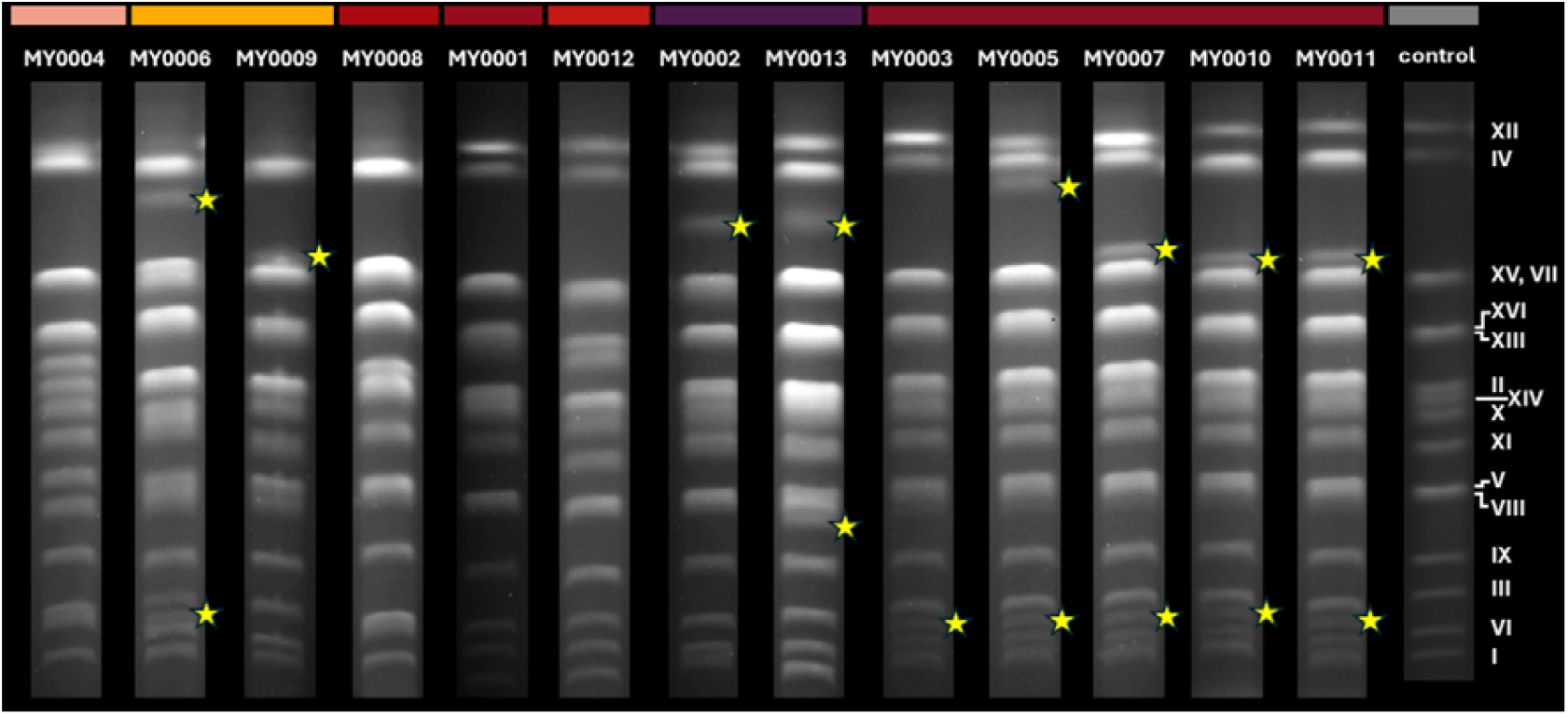
Electrophoretic karyotypes of the mead yeasts, along with a control *S. cerevisiae* sample. Chromosome bands are marked on the right side. Samples are ordered according to clade. Yellow star signs represent supernumerary bands.

### Phenotyping of mead yeasts and experimental fermentation

Several phenotypic characteristics of the mead yeast isolates were compared. All samples produced smooth or relatively smooth whitish colonies on YPD agar, MY0005 and MY0007 produced sectors after 1 week of incubation at 22°C on colony edges (Figure 4a-b.). Strong or weak invasivity into agar was observed in the majority of samples (Figure 4c-d.), as only MY0004, MY0006, MY0001, and MY0007 were non-invasive (Table 3.). Thus, among the five COW clade samples, invasivity was variable. Sporulation occurred in all samples, but MY0012 of the WEW clade and the aforementioned MY0007 produced only deformed asci. Flocculation or foam formation was not observed among the samples, neither in YPD liquid medium, nor during the experimental mead fermentation that was carried out in triplicates with all samples. During the experimental fermentations, conspicuous haziness was observed in meads fermented with MY0004 and MY0006. These samples had a deeper yellow hue as well (Figure 4e-f.).

**Table 3.**
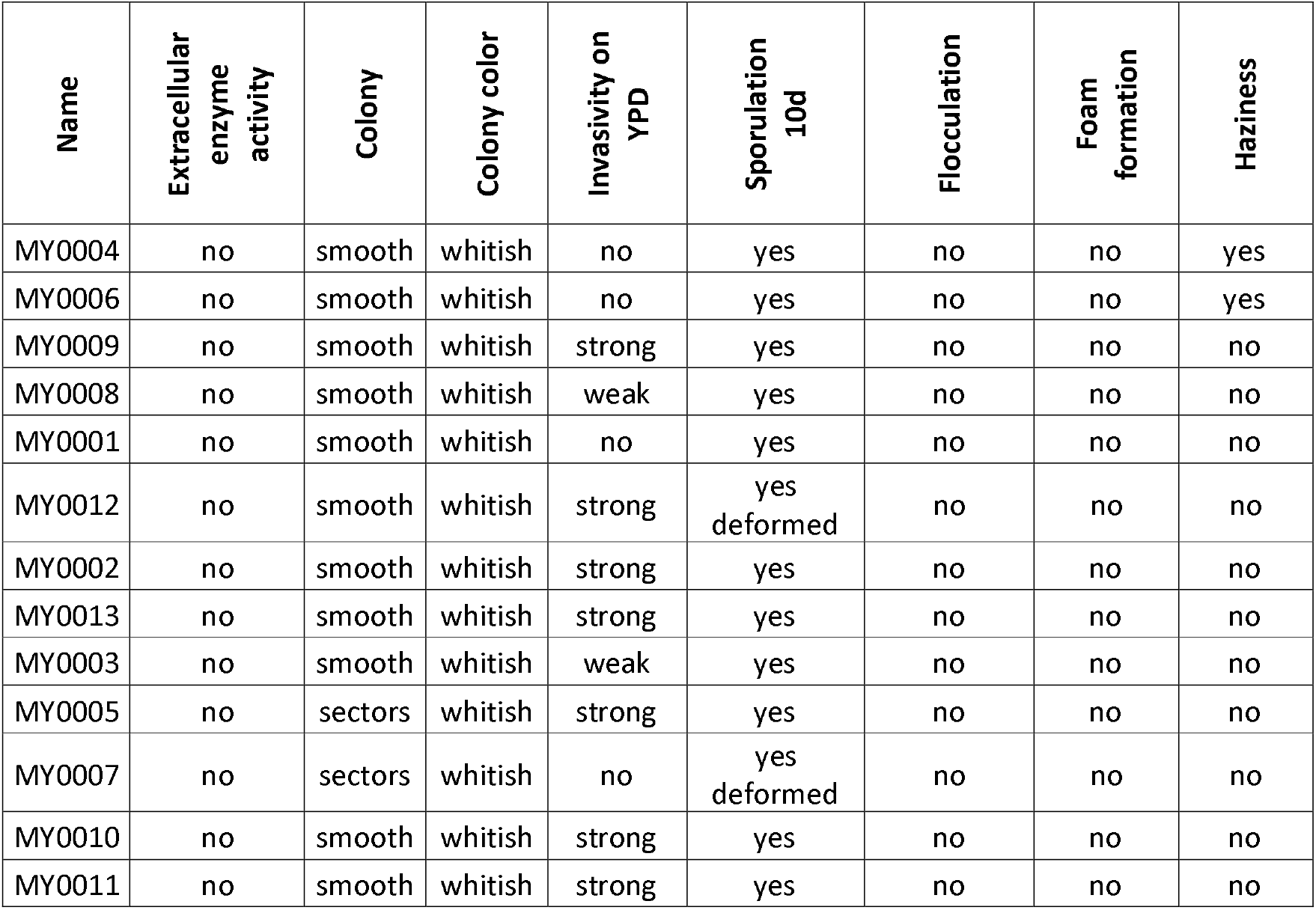
Phenotyping results on the 13 mead yeasts. Samples are ordered according to clade.

**Figure 4.**
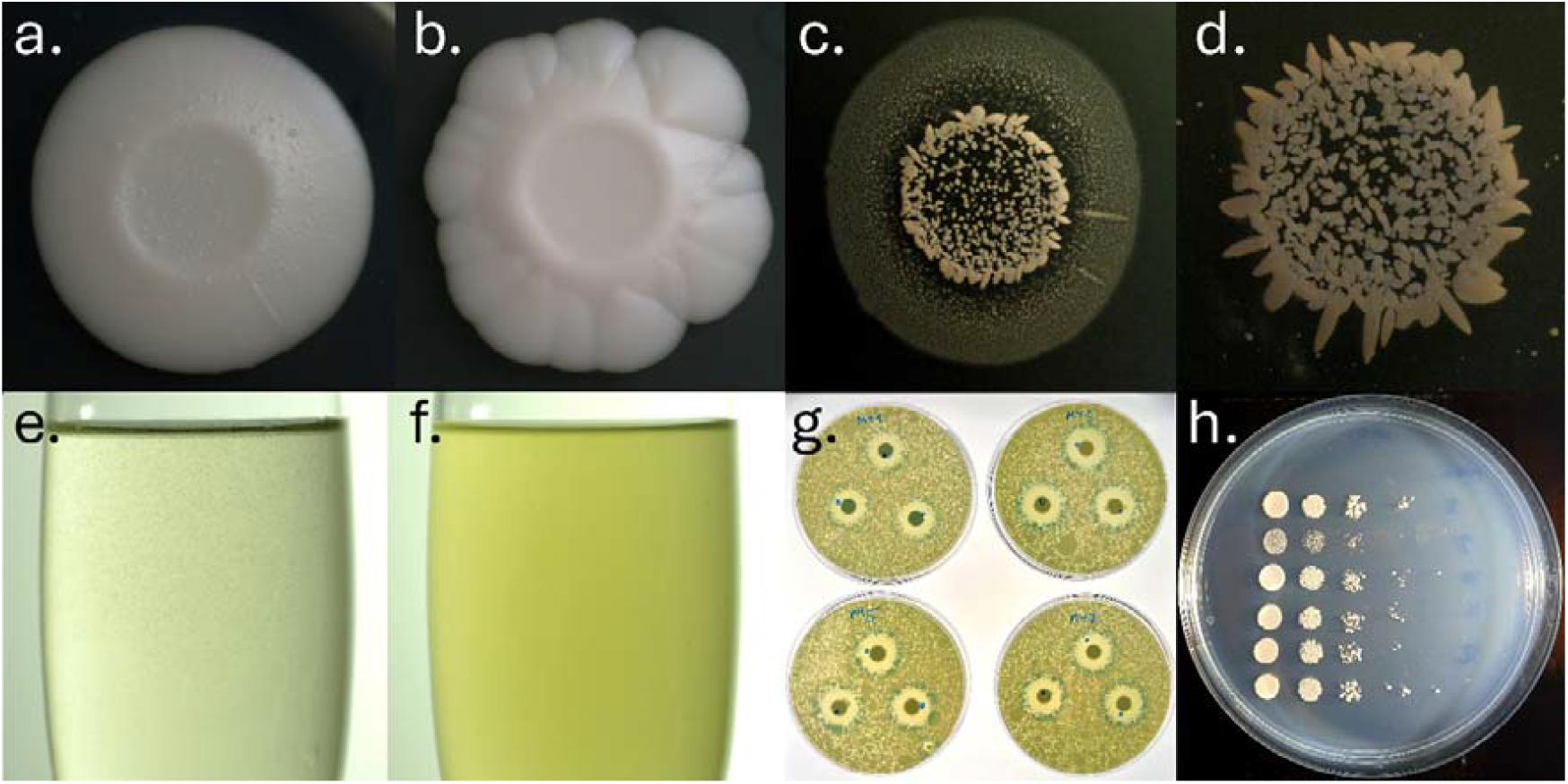
Examples of phenotyping results. Experiments were performed with students during the course of the laboratory class. a. Colony morphology of MY0001. b. Colony morphology of MY0007. c. Invasivity into agar in the case of MY0002. d. Invasivity of MY0004. e. Mead sample fermented with MY0001. f. Mead sample fermented with MY0004. g. Killer activity tests of the mead yeasts. h. Spot plate phenotyping of various mead yeasts (MY0008–MY00013) on SD medium containing only 10% of the usual ammonium sulfate amount (partial nitrogen starvation).

Killer activity and sensitivity to killer toxins was tested for all mead yeasts, and only MY0001 (of the AUW3 clade), and the COW clade MY0003, MY0005, and MY0007 showed killer activity (Figure 4g.). Thus, among the five COW samples, MY0010 and MY0011 have lost their killer phenotype (Figure 5a.). Sensitivity to K1 toxin was observed for all samples (Figure 5b.) and the most sensitive yeasts were MY0012, MY0010, and MY0011. To K2 toxin, only MY0004, MY0006, MY0009, MY0008, MY0012, and MY0002 showed sensitivity, thus, none of the COW, AUW1, or AUW3 clade mead yeasts (Figure 5c.).

**Figure 5.**
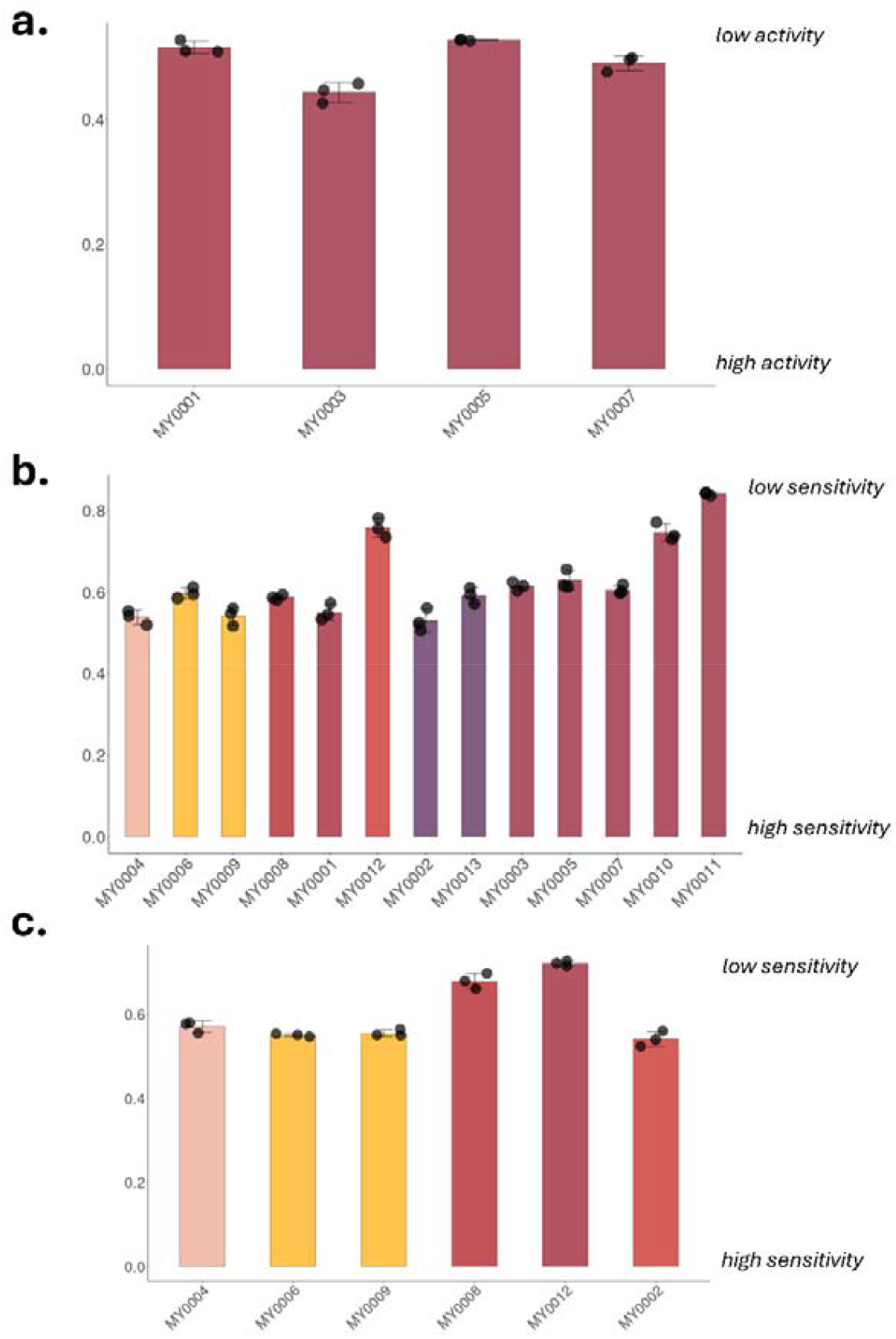
Results of killer sensitivity and activity tests. Values on axis *y* show zone diameter divided by colony diameter values. **a.**Killer activity observed for four samples, all in the COW clade. **b**. Sensitivity to K1 toxin. **c**. Sensitivity to K2 toxin.

Following the general phenotyping of the mead yeasts, we tested their stress tolerance via the spot plate method. Mead yeasts were tested under high temperature, glucose and nitrogen starvation, vitamin starvation. Furthermore, their survival in various concentrations of ethanol and growth in high glucose osmotic stress in liquid medium, simulating very high gravity mead fermentations, was also tested. These latter experiments were carried out after pre-adaptation of the yeasts during a simulated high-sugar fermentation. Mead yeasts could tolerate high temperatures (37°C) and all grew well on YPD agar medium. On SD medium, MY0008 (UKB clade) showed slower growth than other samples. Very low glucose or ammonium sulfate concentrations in SLG and SLAD, respectively, often severely diminished growth of the mead yeasts (in the case of MY0004, MY0008, MY0012, MY0002, MY0013, and MY0003). At concentrations of 0.5 g/l of the nitrogen source, MY0008 and MY0002 showed the most restricted growth, suggesting higher nitrogen requirements (Figure 4h.). Low biotin concentrations severely diminished the growth of MY0008, while low thiamin affected MY0008 and MY0003 the most (Table 4.). The most ethanol tolerant strain was MY0004, capable of surviving at 22% v/v ethanol for up to 7 days. MY0001, MY0012, and MY0010 survived at 18%, 17%, and 17%, respectively, the other samples showed survival between 15% and 16% (Table 4.). High glucose osmotic stress was least tolerated by MY0008, which did not grow or ferment above 30% w/v. The other mead yeasts showed slow growth and fermentation at up to 50% w/v, but not at 60% w/v (Table 4.).

**Table 4.** Stress phenotyping of the mead yeasts. Yeasts were subjected to high temperature and to starvation conditions on agar media, and were tested in liquid media for ethanol tolerance (survival for 7 d) and for high osmotic stress in simulated fermentations.

|  | Growth score |  |  |  |  |  |  |  |  |  | Survival | Growth in high glucose (w/v %) 22°C |  |  |  |  |
| --- | --- | --- | --- | --- | --- | --- | --- | --- | --- | --- | --- | --- | --- | --- | --- | --- |
| Name | SD 22°C | SD 30°C | SD 37°C | YPD 22°C | SLG 22°C | SLAD 22°C | Nitrogen 5% 22°C | Nitrogen 10% 22°C | Low Biotin 22°C | Low Thiamin 22°C | Ethanol 22°C | 2% | 30% | 40% | 50% | 60% |
| MY0004 | 3 | 4 | 4 | 4 | 1 | 1 | 2 | 4 | 2 | 2 | 22% | + | + | + | +- | - |
| MY0006 | 3 | 4 | 4 | 4 | 2 | 2 | 2 | 4 | 2 | 2 | 15% | + | + | + | +- | - |
| MY0009 | 3 | 4 | 4 | 4 | 2 | 2 | 2 | 4 | 2 | 2 | 15% | + | + | + | +- | - |
| MY0008 | 2 | 2 | 2 | 4 | 0 | 0 | 1 | 2 | 1 | 1 | 15% | + | + | - | - | - |
| MY0001 | 3 | 4 | 4 | 4 | 3 | 3 | 2 | 4 | 2 | 2 | 18% | + | + | + | +- | - |
| MY0012 | 3 | 4 | 2 | 4 | 0 | 0 | 2 | 4 | 2 | 2 | 17% | + | + | + | - | - |
| MY0002 | 3 | 4 | 4 | 4 | 2 | 1 | 1 | 4 | 2 | 2 | 16% | + | + | + | +- | - |
| MY0013 | 3 | 4 | 4 | 4 | 0 | 0 | 2 | 4 | 2 | 2 | 15% | + | + | + | +- | - |
| MY0003 | 3 | 4 | 4 | 4 | 1 | 0 | 2 | 4 | 2 | 1 | 15% | + | + | + | +- | - |
| MY0005 | 3 | 4 | 4 | 4 | 3 | 3 | 2 | 4 | 2 | 2 | 16% | + | + | + | +- | - |
| MY0007 | 3 | 4 | 4 | 4 | 2 | 2 | 2 | 4 | 2 | 2 | 15% | + | + | + | +- | - |
| MY0010 | 3 | 4 | 4 | 4 | 2 | 3 | 2 | 4 | 2 | 2 | 17% | + | + | + | +- | - |
| MY0011 | 3 | 4 | 4 | 4 | 2 | 2 | 2 | 4 | 2 | 2 | 16% | + | + | + | +- | - |

### Analytics and characterization of experimental mead fermentations

As mead yeasts are marketed for fermenting honey must, we tested their performance in small-batch fermentations in triplicates, enabling us to compare oenological characteristics and aroma compounds of the meads. After fermentation was completed, the mead samples had densities between 0.993 and 1.016 g/cm^3^, with the highest values observed in the case of MY0004 and MY0008 (Figure 6a., Supplementary File S2.). Regarding ethanol content, only MY0004 and MY0008 showed relatively high standard deviation among the triplicate mead samples. Among all yeasts, ethanol content reached 11.05 to 14.3% v/v. All mead yeasts except the former two sweet mead yeasts reached ethanol levels of at least 13.4% (Figure 6b., Supplementary File S2.). Similar observations were made on extract content, where the two aforementioned samples showed highly different, higher values than the other yeasts, and were more variable among the triplicate samples. They also had in general higher glucose amounts remaining after fermentation (Figure 6c-d., Supplementary File S2.). These mead yeasts were also different from the 11 other mead yeasts when remaining fructose content was measured (Figure 6e., Supplementary File S2.), and showed high and highly variable remaining fructose in the meads. Glycerol content was less variable, despite some significant differences among samples (Figure 6f., Supplementary File S2.), reaching levels of 7.6 to 9.2 g/l. While the pH of the meads was less variable (ranged from 3.13 to 3.46), the titratable acidity values, most important for sensory perception, showed larger differences. MY0002, MY0003, and MY0004 all had values of less than 2 g/l when titrating to pH 7.0, and, along with MY0008, also had the lowest values when titrating to pH 8.2 (Figure 6g-i., Supplementary File S2.). Tartaric acid production was not merely variable among clades, but also variable among members of the COW clade with 0.65 to 1.47 g/l values measured (Figure 6j., Supplementary File S2.). Total volatile acids were highest in MY0012, MY0001, and MY0013, while total polyphenol content was highest in the case of MY0005, with a marked variability in the COW clade (Figure 6k-l., Supplementary File S2.). In this clade, meads fermented with MY0003 had an undetectable level of polyphenols, while other meads had 0.1 to 0.66 g/l values for polyphenol content (Figure 6l., Supplementary File S2.).

**Figure 6.**
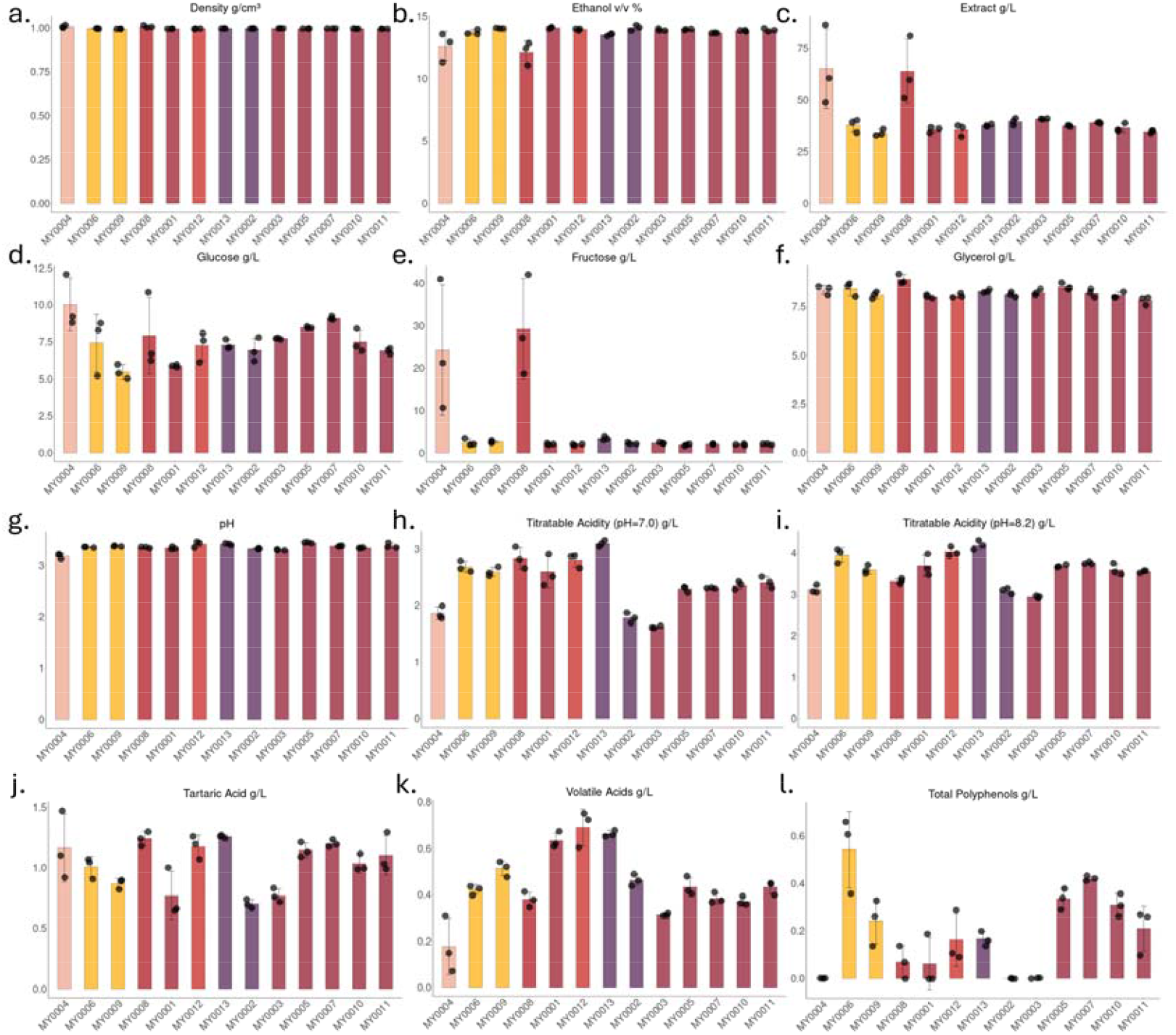
Results on oenological measurements conducted on meads fermented with various yeasts. Means and individual measurements for triplicate samples are shown for each yeast. Bar charts are colored according to clade. **a.** Densities of meads. **b**. Ethanol content. **c**. Extract content. **d**. Remaining glucose after fermentation. **e**. Remaining fructose after fermentation. **f**. Glycerol content. **g**. pH values of the meads. **h**. Titratable acidity to pH 7.0. **i**. Titratable acidity to pH 8.2. **j**. Tartaric acid content. **k**. Volatile acid content. **l**. Total polyphenol content of the experimental meads.

In addition to oenological characterization of the mead samples, we also measured altogether 42 individual volatile metabolites with HS-GC/MS (Supplementary Table S2). Of these, 9 remained unidentified. We applied a semi-quantitative method to analyze differences in the triplicate mead samples fermented with the 13 yeasts, where the area under curve values were expressed as a fraction of the highest value among all samples’ all metabolites. The various yeasts displayed prominent differences in their metabolite profiles (Figures 7-8 and Supplementary Table S2). In most samples, ethyl acetate (fruity notes, Pubchem, accessed 19^th^ February 2026) was among the metabolites with the highest relative values and the highest producer was MY0003, with significant differences when compared to 9 of the 12 other yeasts). Across all mead samples, ethyl caprylate (ethyl octanoate; fruity, sweet, waxy, pineapple notes, Pubchem, accessed 19^th^ February 2026) was on average the second-highest produced metabolite. MY0001, MY0008, MY0012, and MY0013 produced notably smaller amount than the rest of the samples (Figures 7,8a and Supplementary Table S2). Isoamyl alcohol (fruity notes in lower concentrations, Pubchem, accessed 19^th^ February 2026) production was detected in all meads. This metabolite was on average the 3^rd^ among all in relative concentrations, with significant differences among several yeasts (Supplementary Table S2). Notable other differences in the meads include high ethyl caprate (ethyl decanoate, waxy, fruity, sweet apple notes, Pubchem, accessed 19^th^ February 2026) production in the case of MY0004 and very low production in the case of MY0008, MY0012, and MY0013. Among the 12 most prominent metabolites, all showed significant differences in relative values (one-way ANOVA or Kruskal-Wallis, Supplementary Table S2) when all samples were compared (Supplementary Table S2). While most of the clades of the species *S. cerevisiae* with mead yeast samples were only represented by one or two samples in our dataset (Figure 7), the Commercial Wine clade (Figure 8a) and its 5 individual samples allows for more detailed comparisons. The metabolite profiles of these yeasts were remarkably similar, with ethyl acetate as the most pronounced compound, followed by ethyl caprylate (lower levels detected for MY0003), isoamyl alcohol, an unidentified metabolite, and ethyl caprate. A principal component analysis taking all measured metabolites into account clustered the Commercial Wine clade samples near each other (Figure 8b), along with MY0002, MY0004, and MY0006. Additionally, MY0008, MY0009, MY0012 were clustered together, while MY0001, a relatively high aroma producer, and MY00013, the yeast with overall lowest aroma production, were placed separately from other samples and from each other in the analysis. Notably, mead fermented with MY0006 displayed relatively high levels of a metabolite identified as 2-undeca-2,5-diynoxyoxane, an eicosanoid substance not found in the case of other mead yeasts. Its sensory characteristics are unknown (Pubchem, accessed 19^th^ February 2026).

**Figure 7.**
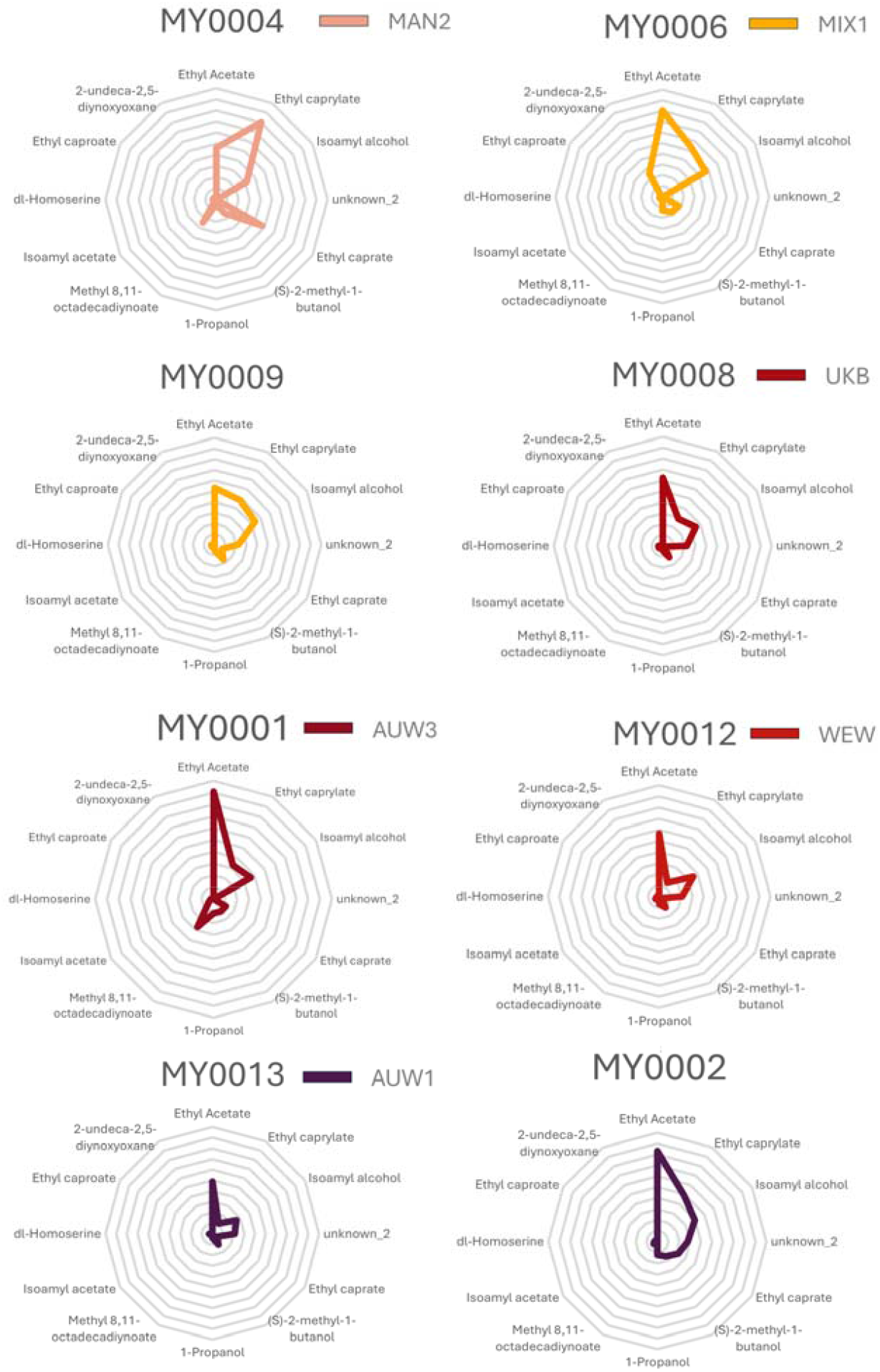
Results of HS-GC/MS measurements for the meads fermented with yeasts of the MAN2, MIX1, UKB, AUW3, and AUW1 clades. Each radar chart is scaled from 0 to 1 (innermost point and outermost circle, respectively). Values show semi-quantitative measurements of each identified compound, expressed as a fraction of the highest measured area under curve value (value of 1; among all replicates of all mead yeasts). Clades are color-coded, mead yeast identifiers are on top of each radar chart. Only the 12 metabolites with highest average values among all replicates of all mead yeasts are shown, continuous lines show averages of triplicate fermentations.

**Figure 8.**
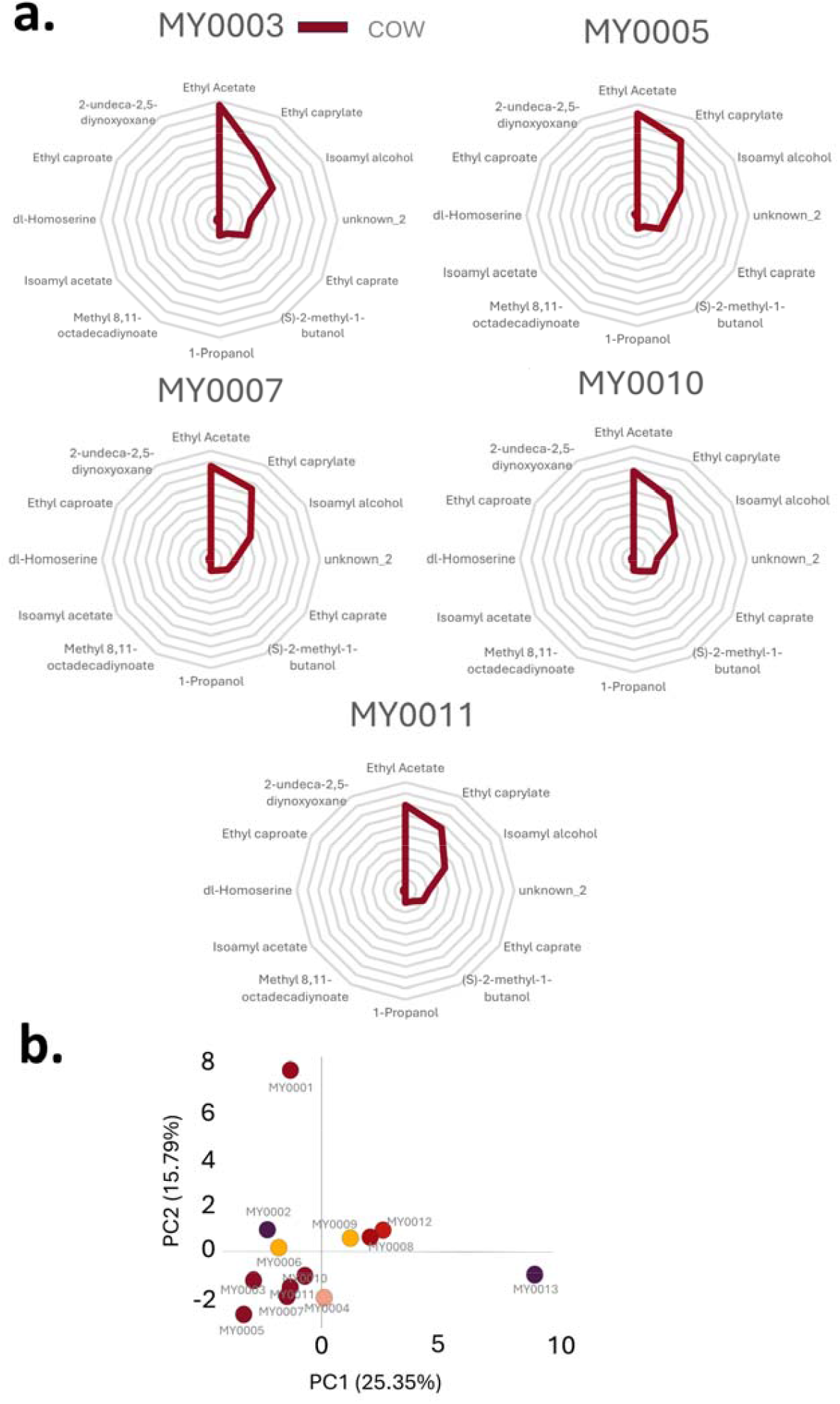
**a.** Results of HS-GC/MS measurements for the meads fermented with yeasts of the COW (Commercial wine) clade. Each radar chart is scaled from 0 to 1 (innermost point and outermost circle, respectively). Values show semi-quantitative measurements of each identified compound, expressed as a fraction of the highest measured area under curve value (value of 1; among all replicates of all mead yeasts). Clades are color-coded, mead yeast identifiers are on top of each radar chart. Only the 12 metabolites with highest average values among all replicates of all mead yeasts are shown, continuous lines show averages of triplicate fermentations. b. Principal component analysis based on the 42 individual metabolites’ scaled semi-quantitative measurements identified in HS-GC/MS analysis (Supplementary Table S2). Scaling was performed as described for Figure 7 and 8a. PCA analysis was based on averages for each metabolite from the triplicate fermentations. Spots are color-coded for clades. PC1 (*x* axis) and PC2 (*y* axis) shown.

## Discussion

In the present work, we characterized a group of commercially available *Saccharomyces* yeasts that have not been scientifically evaluated and compared before, namely the mead yeasts. Although mead is an ancient fermented beverage with a growing number of scientific articles focusing on its characteristics, yeasts’ role in the quality and taste of honey-wine is rarely researched, as mead is mostly considered a specialty product with only small-scale production especially compared to wine or beer. Yet, its obscurity provides an opportunity as a novel research project partially integrated into a microbiological lab class. To compare the mead yeasts, we obtained 13 products from various countries and subjected them to detailed phenotypic tests, genetic characterization, and made experimental fermentations with them.

First, we compared the genomes of the samples with the few available genomes of yeasts isolated from traditional mead fermentations. Ethiopian T’ej isolates were found to cluster with the African Beer yeasts (Figure 2) and with the Mixed Origins 1 clade, the latter likely represent recently repurposed commercial baker’s yeasts. These latter were collected in the 2010s. The African Beer clade isolates are much more likely to be naturally occurring locally in honey must fermentations, especially as they were also isolated from geshu leaves used for flavoring T’ej, and as they have been identified in collections from before the 1960s and from the 2010s as well. Historic T’ej was, however, not uniform in terms of *Saccharomyces* clades, as two samples from before the 1960s clustered with some genomes of the Australia Wine 1 clade.

There is a very small number of isolates available from mead fermentations in Europe from the 20^th^ century, among these, two samples fell into the Georgia Wine clade, while one into the Mixed Origins 1 clade. The currently marketed mead yeasts, on the other hand, showed diverse origins and none of them clustered with the African Beer or Georgia Wine yeasts.

Based on their relatedness to other commercially available yeasts, they are likely relatively recently repurposed yeasts from fermentation environments such as liquor making (Huangjiu clade, MY0004), ale brewing (MY0008, UK Beer clade), or wine and Champagne making (various wine clades). The origin of the two Mixed Origins 1 samples, MY0006 and MY0009, is complicated to evaluate. In contrast to the aforementioned Ethiopian samples in Mixed Origins 1, these two yeasts do not show very close relatedness to globally available active dry baker’s yeast products and fall into a smaller group of strains.

Despite originating from various countries and manufacturers, 5 of the 13 mead starter product samples were found to be very closely related to each other and to globally available wine and Champagne yeast starter strains in the Commercial Wine clade (Table 1, Figures 1-2). Regardless of their close relatedness, these yeasts differed in genome structural variations (Table 2., Supplementary File S1, Figure 3).

Phenotyping revealed that none of the samples were strong flocculators and two were unable to form proper asci during sporulation experiments (MY0007 and MY0013). Variability in invasive growth on agar medium was also observed (Table 3.). The MY0004 and MY0006 produced pronounced haziness in honey must fermentation, the other 11 yeasts produced clear and lightly colored meads. Killer activity was found in MY0001 and three yeasts of the Commercial Wine clade. Other yeasts of this clade are also often marketed as killer yeast starters. Thus, two yeasts in the latter clade have apparently lost their killer phenotype. All yeasts were sensitive to K1 killer toxin, and only six to K2 toxin (Figure 5). Yeasts tested here also showed differences in their stress tolerance and nitrogen and vitamin requirements (Table 4). Together with the killer phenotype and haziness, these results have practical considerations during meadmaking. None of the samples showed good fermentation in 50% w/v glucose, thus they may not be suitable for more extreme mead fermentations. Ethanol tolerance differed substantially among the samples, with MY0004 (22%) standing out. The second-most tolerant yeast was MY0001, with 18%. Six samples had a maximum ethanol tolerance of 15%, placing some limitations on the meads fermented with these (Table 4). Surprisingly, all of the tested yeasts except for MY0012 grew well even at 37°C under laboratory conditions, although their normal fermentation environment is characterized by lower temperatures. In a recent work, one human isolate from the Commercial Wine clade from the clinics of the University of Debrecen was described, with close relatedness to the five mead yeasts of the COW clade and to wine strains EC 1118, CBC-1, Fermicru 4F9, WY4766 (Harmath et al., 2026). According to our current results, these yeasts are commonly able to withstand the human body temperature. In experimental fermentations of honey must, conducted in triplicates, the individual yeasts showed several differences. Notable is the lower and more variable ethanol production in the case of MY0004 and MY0008 (Figure 6.) compared to other yeasts, especially as the former was the most ethanol-tolerant among all yeasts. These samples had higher extract and higher remaining fructose content. Total polyphenol content of the experimental meads, was highly variable, along with titratable acidity, volatile acid production, and tartaric acid content, while glycerol production showed less variability (Figure 6., Supplementary Table S1.). The aroma compounds as measured by HS-GC/MS clearly differentiated MY0001 and MY0013 from the other samples, while the rest of the yeasts formed two clusters based on the identified metabolites and their relative amounts in our semi-quantitative analysis (Figures 7-8.). Metabolites associated with fruity notes were prominent among the identified molecules (Supplementary Table S2). In accordance with the variable oenological characteristics and metabolite profiles of the 13 mead yeasts, the meads fermented with them differed in their sensory evaluation during tasting, with sweet mead yeasts (MY0004 and MY0008) producing meads perceived mostly as sweet, fruity, while fruitiness was also prominent in MY0001, along with vinous notes. Notably, the mead with MY0013, the lowest aroma producer, was perceived as the most neutral, while MY0006 as perceived as bitter, complex in taste, and mostly neutral in smell (FigShare, https://figshare.com/projects/Comparison_of_mead_yeasts/271906).

Our results thus showed that the currently available mead yeast starter strains belong to various clades of the *S. cerevisiae* species, and they differ from the few available historic and traditional mead yeasts in culture collections. In line with their diverse origins, they are different in several key phenotypic characteristics, oenological traits and in aroma production, and in killer activity. Mead yeasts, as a collection of microbes from a not-yet researched source proved to represent an engaging opportunity to discuss applied microbiological topics in a university class setting according to our experience. It allowed designing and performing experiments at a BSc-level laboratory class, amended with more detailed analyses on genomics and analytics subsequently.

## Data availability

Raw sequencing reads used in this study are deposited in NCBI SRA under BioProject PRJNA1420404. Raw data for killer tests and sensory evaluation of meads is uploaded to FigShare (https://figshare.com/projects/Comparison_of_mead_yeasts/271906).

## Acknowledgements

The research was funded by the National Research, Development and Innovation Office (NKFIH FK 138910 to W.P.P.). This project received funding from the NKFIH 2020-1.1.2-PIACI-KFI-2020-00130 grant (to M.S.). This project has received funding from the HUN-REN Hungarian Research Network (to I.P.).

## Author Contributions

Conceptualization: W.P.P, Z.K.; methodology: W.P.P., Z.K., B.N., G.H., A.T.; formal analysis: W.P.P., Z.K., B.N.; A.T., G.H., A.H., Zs.A., University of Debrecen Biotechnology BSc class of 2026; resources, W.P.P., I.P., M.S.; data curation, W.P.P., B.N.; writing—original draft preparation: W.P.P., A.T.; writing—review and editing: W.P.P., B.N.; visualization, W.P.P., B.N.; supervision: W.P.P.; B.N., A.H.; project administration: W.P.P.; funding acquisition: W.P.P., I.P., M.S.

