## Supplementary File S1 for "A comparison of commercially available *Saccharomyces* mead yeasts"

**Supplementary File S1.** Additional results on comparative genomics of mead yeasts.

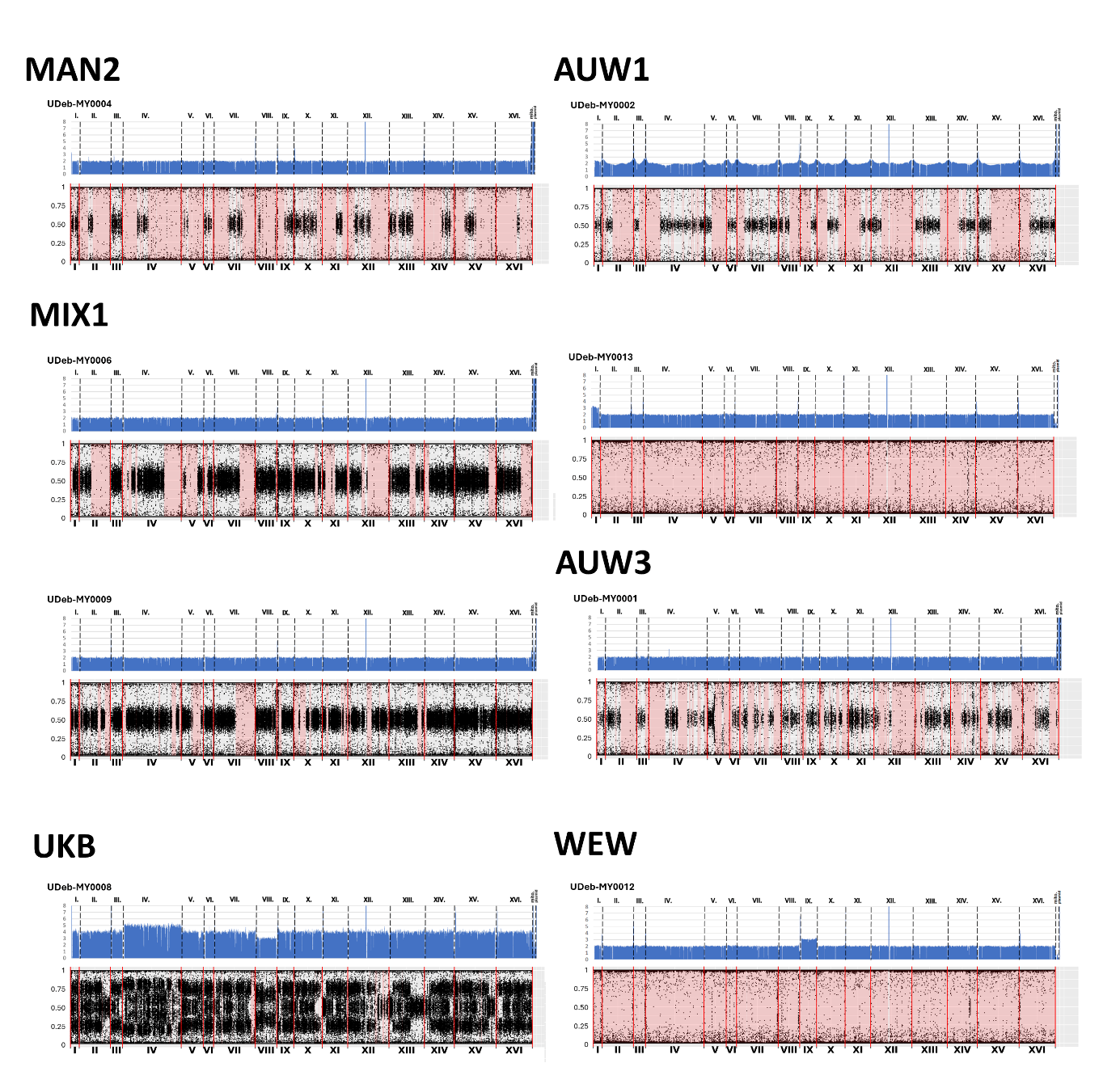

Copy-number corrected coverage plots (top parts of figure pairs) and allele ratio plots (bottom parts) of the 6 mead yeasts in clades MAN2, MIX1, UKB, AUW1, AUW3, and WEW. Figures are ordered according to clade. Called ROH regions are shown in light red background along the allele ratio plots.

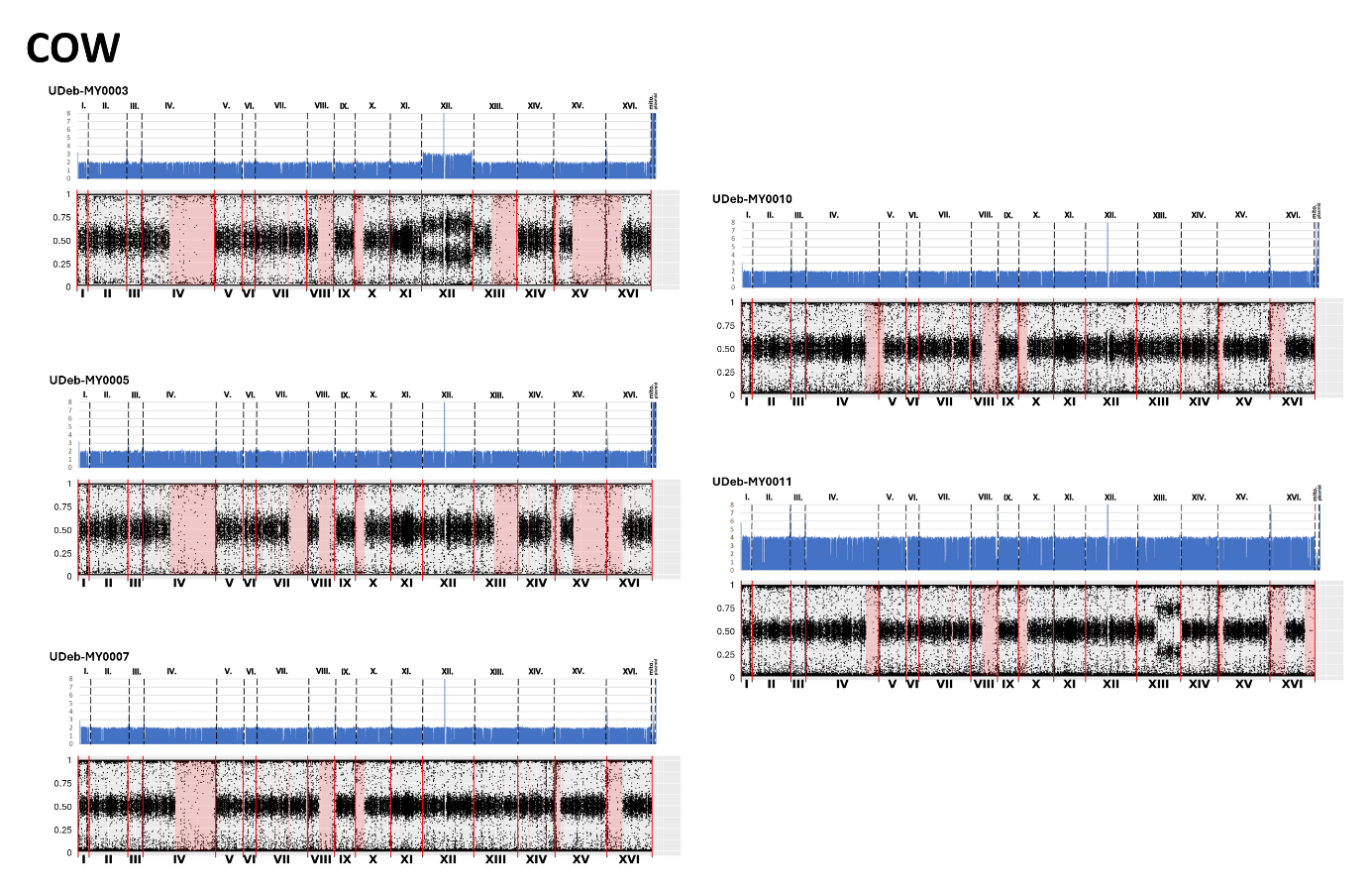

Copy-number corrected coverage plots (top parts of figure pairs) and allele ratio plots (bottom parts) of the 5 mead yeasts in clade COW. Called ROH regions are shown in light red background along the allele ratio plots.
